# A guide to examining intramuscular fat formation and its cellular origin in skeletal muscle

**DOI:** 10.1101/2022.03.06.483182

**Authors:** Connor D. Johnson, Lylybell Y. Zhou, Daniel Kopinke

## Abstract

Fibro-adipogenic progenitors (FAPs) are mesenchymal stromal cells that play a crucial role during skeletal muscle homeostasis and regeneration. FAPs build and maintain the extracellular matrix that acts as a molecular myofiber scaffold. In addition, FAPs are indispensable for myofiber regeneration as they secrete a multitude of beneficial factors sensed by the muscle stem cells (MuSCs). In diseased states, however, FAPs are the cellular origin of intramuscular fat and fibrotic scar tissue. This fatty fibrosis is a hallmark of sarcopenia and neuromuscular diseases, such as Duchenne Muscular Dystrophy. One significant barrier in determining why and how FAPs differentiate into intramuscular fat is effective preservation of adipocytes, especially in frozen tissue sections. Conventional methods of skeletal muscle tissue processing, such as snap-freezing, do not properly preserve the morphology of individual adipocytes, thereby preventing accurate visualization and quantification. Here, we describe a protocol that provides robust preservation of adipocyte morphology in skeletal muscle sections allowing visualization, imaging, and quantification of intramuscular fat. We also outline how to process a portion of muscle tissue for RT-qPCR, enabling users to confirm observed changes in fat formation by viewing differences in expression of adipogenic genes. Additionally, we will describe how our protocol can be adopted to visualize adipocytes by whole mount immunofluorescence of muscle samples. Finally, we will outline how to combine this protocol with genetic lineage tracing of *Pdgfrα*-expressing FAPs to study the adipogenic conversion of FAPs. Our protocol consistently yields high-resolution and morphologically accurate immunofluorescent images of adipocytes that, along with confirmation by RT-qPCR, allows for robust, rigorous, and reproducible visualization and quantification of intramuscular fat. Together, our analysis pipeline is the first step to improve our understanding of how FAPs differentiate into intramuscular fat and provides a framework to validate novel interventions to prevent fat formation.

## INTRODUCTION

The infiltration of healthy muscle tissue with fatty fibrosis is a prominent feature of Duchenne Muscular Dystrophy (DMD) and other neuromuscular diseases, as well as sarcopenia, obesity, and diabetes ^1,2^. Although increased fat infiltration in these conditions is strongly associated with decreased muscle function, our knowledge of why and how intramuscular fat forms is still limited. FAPs are a multipotent mesenchymal stromal cell population present in most adult organs including skeletal muscle ^3–10^. With age and in chronic diseases, however, we and others have found that FAPs produce fibrotic scar tissue and differentiate into adipocytes, which are located between individual myofibers and form intramuscular fat ^7,11–20^.

To start combating intramuscular fat formation, the mechanisms on how FAPs turn into adipocytes need to be defined. PDGFRα is the “gold-standard” marker in the field to identify FAPs within murine and human muscle ^3,6–9,21–24^. As a result, several tamoxifen-inducible Cre lines, under the control of the *Pdgfrα* promotor, have been generated allowing to genetically manipulate FAPs, using the Cre-LoxP system, *in vivo* ^24–26^. For example, by combining this inducible Cre line with a genetic reporter, lineage tracing of FAPs can be performed, a strategy we have successfully applied to fate map FAPs in muscle and white adipose tissue ^9,27^. Besides lineage tracing, these Cre lines provide valuable tools to study the FAP-to-fat conversion.

One major obstacle to define the mechanism of the adipogenic conversion of FAPs into intramuscular fat is the ability to rigorously and reproducibly quantify the amount of intramuscular fat that has formed under different conditions. The key is to balance preservation of muscle and fat tissue, and match this with the available staining methods to visualize adipocytes. For example, skeletal muscle is often snap-frozen without prior fixation, which preserves myofibers but disrupts adipocyte morphology (**Figure 1**). In contrast, fixation followed by paraffin embedding, while displaying the best tissue histology, including that of adipocytes, removes all lipids, thereby rendering most lipophilic dyes, such as the commonly used dye Oil Red O, unusable.

**Figure 1.**
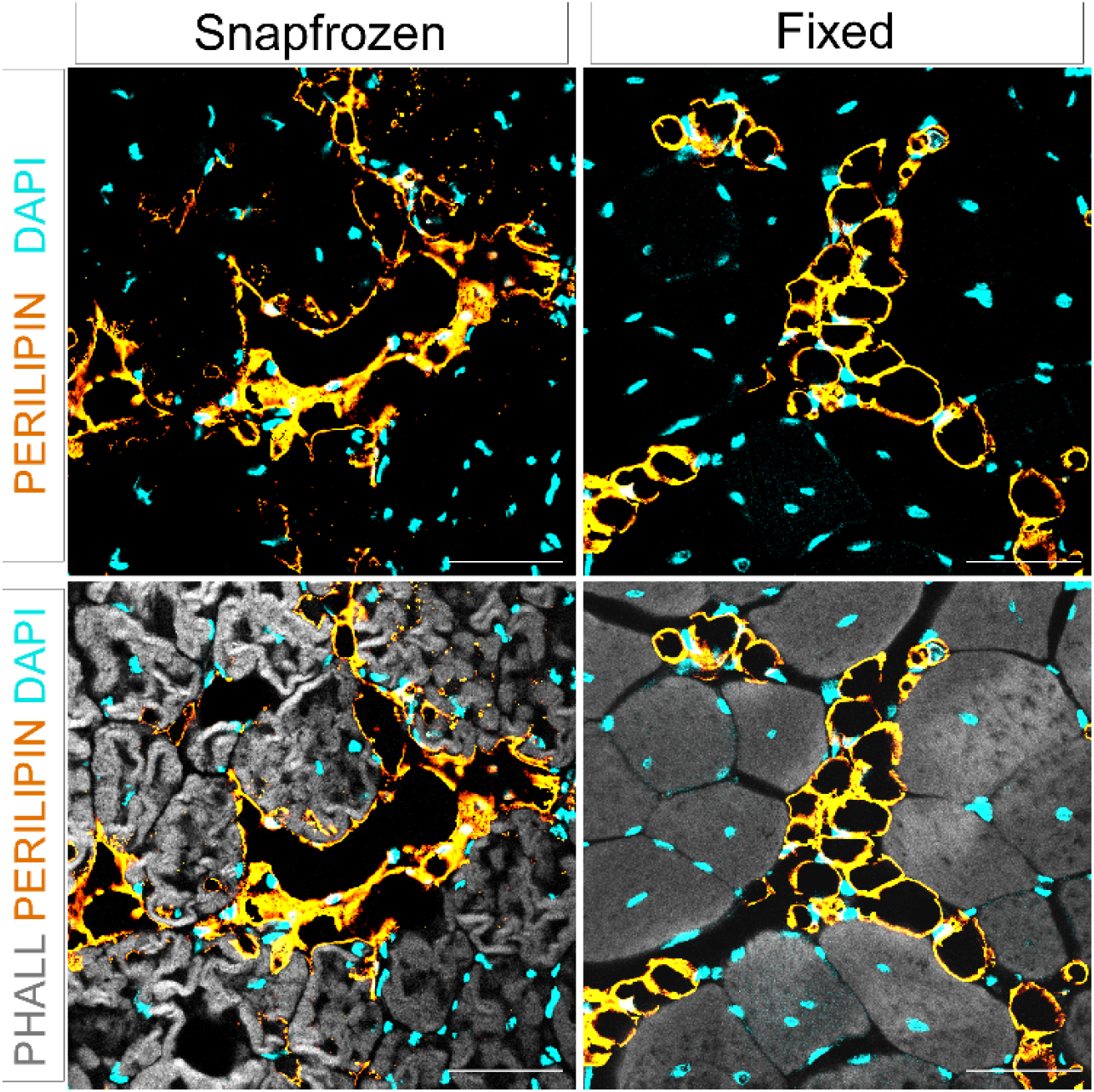
Intramuscular fat in snap-frozen versus fixed muscle tissues. Immunofluorescent images showing adipocytes (yellow), myofibers (gray), and nuclei (cyan) within both snap-frozen and fixed TAs at 21 days post Glycerol injury. Scale bars: 50 μm.

Here, we describe a protocol (**Figure 2**) which preserves myofiber and adipocyte morphology, and allows visualization, and analysis, of multiple cell types. Our approach is based on immunofluorescence staining of adipocytes in paraformaldehyde (PFA)-fixed muscle tissue, which allows for co-staining with multiple antibodies. It can also be easily adapted to spatially display intramuscular fat in intact tissue using whole mount imaging, thereby providing extensive information on the cellular microenvironment of fat within muscle. In addition, our protocol can be combined with our recently published approach to determine the cross-sectional area of myofibers in fixed muscle tissues ^28^, an important measurement to assess muscle health. We also outline how to combine this approach with genetic lineage tracing to fate map the differentiation of FAPs into adipocytes. Thus, our versatile protocol enables rigorous and reproducible assessment of FAPs and their differentiation into intramuscular fat in tissue sections as well as in intact tissues.

**Figure 2.**
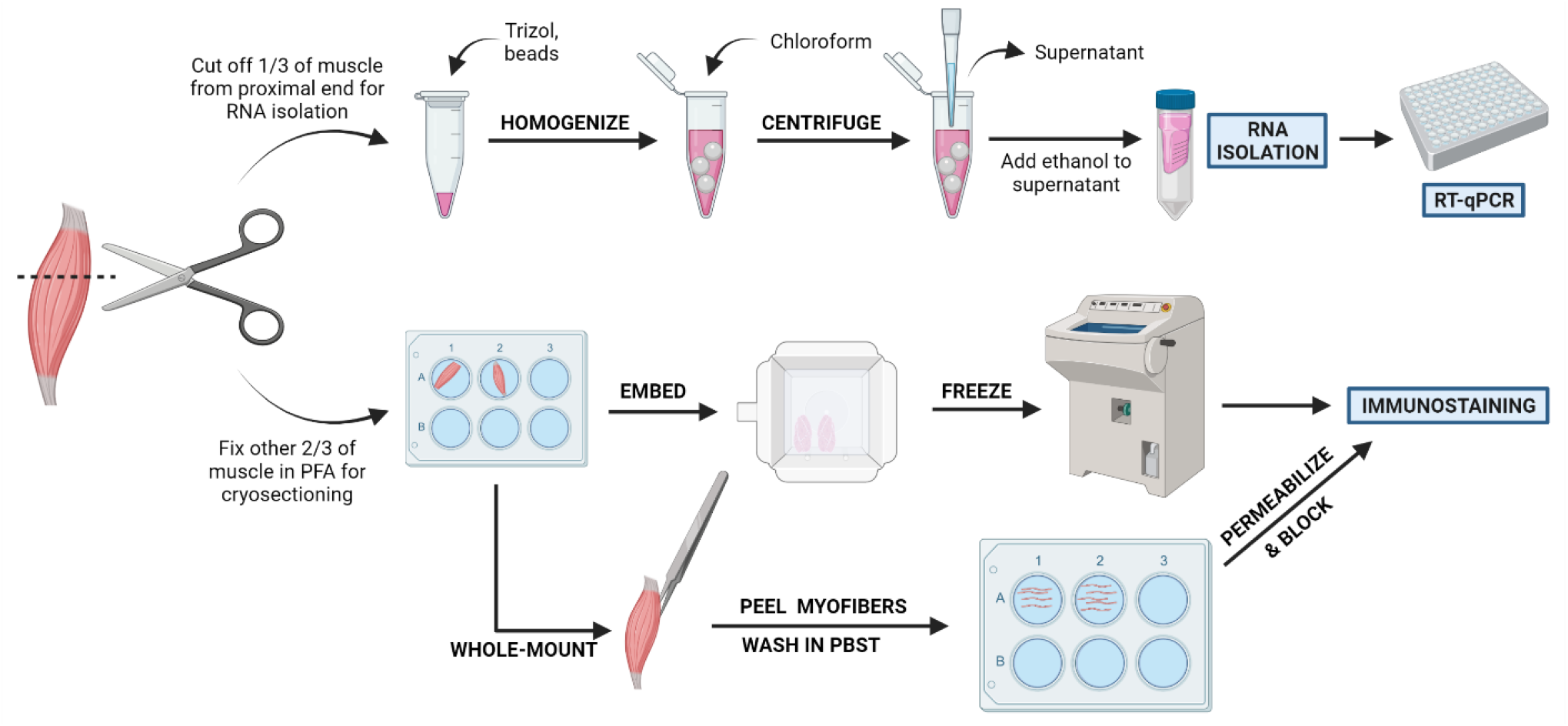
Schematic protocol overview. Schematic overview of tissue processing in which 1/3 of the TA is removed, snap-frozen, and homogenized for subsequent RNA isolation and transcription analysis via RT-qPCR. The other 2/3 of the TA is PFA-fixed and processed for immunostaining on frozen sections or whole-mount fibers.

## PROTOCOL

All animal protocols were approved by the Institutional Animal Care and Use Committee (IACUC) of the University of Florida.

1. **Genetic lineage tracing of FAPs**

NOTE: If genetic lineage tracing of FAPs is not desired, step 1 can be skipped.

1.1 To perform lineage tracing of FAPs, the necessary mouse alleles need to be obtained. For example, several tamoxifen-inducible Cre lines, under the control of the *Pdgfrα* promotor, have been generated to successfully target FAPs including from the Hogan, Rando and Rossi laboratories ^24–26^. As a genetic reporter of Cre activity, several Rosa26 reporter alleles are available such as *Rosa26^EYFP^* ^29^.

1.2. **Tamoxifen Administration Through Oral Gavaging**

CAUTION: Tamoxifen is a carcinogen and should be handled very carefully. Always wear gloves when handling and wear a mask when weighing it as a powder, as there is a danger of inhalation.

1.2.1. Clean the area according to protocol and attach a gavaging needle to a 1 mL insulin syringe (without needle). Draw up 200 μL of Tamoxifen (40 mg/ml) into the syringe.

1.2.2. Scruff *Pdgfrα^CreERT2^; Rosa26^EYFP^*mice by placing them on a flat surface and firmly gripping the base of the tail. Use your free hand to grasp the middle of the mouse with your thumb and index finger, then gently and with slight pressure slide the grip until just past the shoulders. Pinch the skin back with the thumb and index finger, pick up mouse and flip hand so that the mouse is facing you, and tuck the tail in between the pinky and ring finger of the hand holding the mouse.

1.2.3. At this point, the mouse should be well immobilized and unable to move head or arms. Insert the gavaging needle into the mouth and use it to slightly tilt the head of the mouse back, this allows the esophagus to be better accessible.

1.2.4. Carefully and slowly insert the needle into the esophagus, do not force the needle if any resistance is met, the needle should slide down easily. Slowly inject the Tamoxifen. Monitor the mice for 15-20 min to ensure no problems occurred while gavaging.

NOTE: We find that administration of Tamoxifen on two consecutive days will result in ~75-85% recombination efficiency of FAPs without causing any adverse effects. We recommend waiting for 1-2 weeks before inducing injury, which will allow the remaining tamoxifen to be removed from the system and any remaining protein to be turned over.

**2. Injury of Tibialis Anterior Muscle**

2.2. Prepare the anesthetic machine by adding Isoflurane and making sure the tubes to both the mouse chamber and nose cone are open. Clean the chamber and work area with either 70% ethanol or peroxide solution (depending on protocols in place).

2.3. Set the flow oxygen flow rate to 2.5 L/min and the Isoflurane concentration to 2.5%. Add a mouse into the anesthesia chamber and wait ~5 min for it to be anesthetized.

2.4. Perform a toe pinch on the mouse to be injured to ensure they do not respond. Then place the mouse down supine on a clean heating pad and insert nose into nose cone. Again, perform a toe pinch at this time and before injuring to ensure the mouse is fully anesthetized.

2.5. Clean the leg to be injected with a fresh alcohol wipe to disinfect.

2.6. Draw up 30-50 μL of 50% Glycerol (depending on size of mice) into the insulin syringe (with needle). Gently use the side of the needle to brush up the hair on the shin to better see the location of the TA.

NOTE: It is easier to move the hair and achieve better visualization when it is still wet from the alcohol wipe.

2.7. After locating the TA (just lateral to the tibia, it somewhat protrudes through the skin and can be felt with gentle palpation), insert the needle into the TA distally, near the ankle. Fully insert the needle into the muscle and slowly inject the Glycerol while slowly withdrawing the needle, this helps to injure most of the muscle.

NOTE: It is best to insert the needle nearly parallel to the leg, with just a slightly elevated angle. A good injury typically causes a reflex of pulling the foot up to the shin after withdrawing the needle. If the mouse’s toes spread out, you likely hit the extensor digitorum longus (EDL) muscle.

2.8. Place the mouse back in the cage and monitor for about 15-20 min to ensure recovery from the anesthesia.

2.9. Discard the needle in a sharps container and never recap a needle.

**3. Tissue harvest**

3.1. Prepare 4% PFA and place on ice prior to starting harvesting.

3.2. Place any plates (12- or 24-well) being used for muscle fixation on ice and add 4% PFA to each well, making sure that each well has 10-20 times more volume of PFA than the tissue being fixed.

3.3. Sacrifice the mouse according to protocol in place.

3.4. Spray any areas of the mouse to be cut into with 70% ethanol to help keep hair off dissecting area and instruments.

3.5. Begin harvesting any tissues to be used for histology or RNA isolation. Tissues should be snap-frozen or placed in PFA within 10-15 min of sacrificing.

3.6. Spray leg with 70% ethanol

3.7. Use scissors to cut the skin around the top of the leg, near the pelvis (**Figure 3A**).

**Figure 3.**
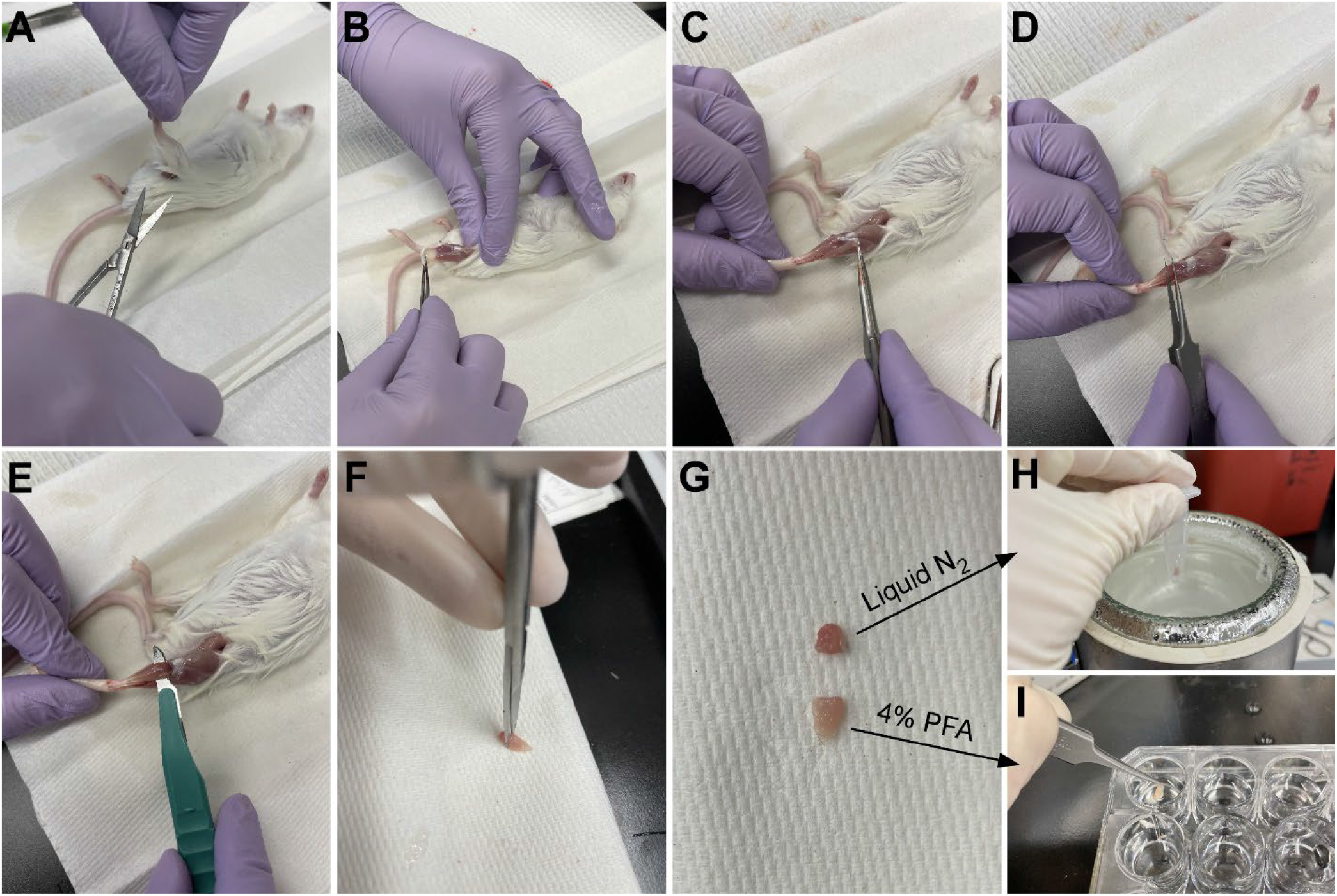
Tissue harvest summary. The skin is cut at the base of the leg (**A**) and the hind limb muscles are exposed (**B)**. Once the epimysium is removed from the TA (**C**), forceps are used to partially separate the muscle and ensure the epimysium has been removed completely (**D**). The TA is cut from the leg with a scalpel and removed after cutting the tendon (**E**). After cutting of the TA into a 1/3 and a 2/3 piece **(G**), 1/3 is snap-frozen in liquid nitrogen for RT-qPCR analysis (**H)** and the other 2/3 is fixed in 4% PFA for histology (**I**).

3.8. Gently pull the skin of the leg from the top down to the ankle (**Figure 3B**).

NOTE: The TA is a teardrop-shaped muscle with a clearly defined tendon attaching to the ankle, it is lateral to the tibia and extends up to the lower knee.

3.9. First, remove the outer connective tissue layer (epimysium) using sharp-tipped tweezers prior to harvesting the TA (**Figure 3C**). A dissecting microscope can be used to better visualize the epimysium.

3.10. Slide the tweezers underneath the TA from the bottom of the muscle, starting at the ankle tendon, and gently pull upwards towards the knee (**Figure 3D**). Stop at the end of the muscle, do not push past the resistance felt at the lower knee.

3.10.2. If there is significant resistance prior to reaching the lower knee, stop and continue removing leftover layers of connective tissue.

NOTE: There is another tendon at the ankle just lateral to the TA tendon, this attaches to the EDL, which is a slender muscle lateral to the TA. Being careful to only slide the tweezers under the TA tendon prevents accidental harvesting of the EDL, but it can also be easily removed after fixation.

3.11. Once the TA has been partially lifted from the leg with the tweezers, use the same motion with a scalpel to sever the connection of the TA to the lower knee (**Figure 3E**). Cut the tendon at the ankle with scissors to fully remove the TA. Only handle the muscle from the tendon to avoid damage to the fibers.

3.12. Cut 1/3 of the TA at the end opposite to the tendon (**Figure 3F**), put in a microcentrifuge tube, and snap-freeze by dropping into liquid nitrogen (**Figure 3H**).

3.13. Submerge the other 2/3 of the tissue in a labelled well with 4% PFA for histology (**Figure 3I**). Be sure to keep track of the time that you put the first and last tissue into fixative. Place on shaker at 4 °C.

3.14. Duration of fixation is dependent on the tissue and its size. While the duration should be determined by each user, we have found that fixing TAs for 2-2.5 h at 4 °C preserves adipocyte morphology very well without causing over-fixation of the tissue.

NOTE: If planning to use TA for whole mount immunofluorescent staining, skip the rest of this protocol until reaching the “Whole Mount Immunofluorescent Staining” section.

3.15. After fixation, remove PFA from the wells and rinse the tissues with cold 1x PBS 2-3 times and then wash 2-3 times with cold 1x PBS for 5 min per wash.

3.16. Remove the PBS from the wells and add enough 30% sucrose in 1x PBS to allow the tissue to float. Place on shaker at 4 °C overnight.

**4. Embedding**

4.1. Prepare the specimen molds to be used for embedding by labelling and filling with enough embedding medium to fully submerge the tissues

4.2. Remove tissues from well, dry off excess sucrose on paper towel and move to medium-filled specimen molds.

NOTE: It is helpful to know in what orientation the mold with be sectioned on the cryostat. This way, you can orient the tissues in the mold in a way that allows the area of interest to be easily accessible. For the TA, this would be achieved by putting the thickest end (opposite from the tendon side) facing the surface that will be sectioned. This allows the TA to be easily sectioned at its thickest area (belly) and allows cross-sectional cutting of the muscle fibers.

4.2.1. Allow the tissues to sit in the embedding medium for ~10 min, this prevents the tissue section from separating from the medium while using the cryostat, which will make the sectioning process easier and more effective.

4.3. Prepare an isopentane slurry by partially submerging a container holding isopentane into liquid nitrogen. There should be enough liquid isopentane in the container to submerge about half of the specimen mold you will be using to embed the tissues.

4.4. Begin freezing the molds by carefully putting them in the isopentane slurry and make sure that about half of the mold is submerged. Also make sure that the mold is freezing equally from all four sides.

4.5. Take the mold out of the isopentane just before the entire mold is visibly frozen over from the top. The amount of time this takes depends on the molds being used.

4.6. Keep the frozen molds in a container with dry ice while freezing the rest of the blocks, then store them at −80° C.

NOTE: Isopentane for freezing can be reused, put into a glass bottle but DO NOT tighten lid until isopentane reaches room temperature. Otherwise, the change in pressure could shatter the bottle.

**5. Sectioning**

5.1. Set cryostat to −22 to −24 °C and add molds containing TAs into cryostat and wait for a minimum of 30 min for temperature acclimatization. In the meantime, label a series of positively charged microscope slides.

5.2. Insert anti-roll plate and align to microtome such that there are minimal nicks in the plate where it makes contact with the specimen block. Secure in place.

5.3. Insert fresh microtome blade in the blade holder and secure in place.

CAUTION: The blade is sharp. When manipulating other parts of the cryostat or frozen molds, cover the blade.

5.4. Remove frozen block from the mold. Add a uniform layer of embedding medium to cryostat chuck and position block in medium. Let sit for 1 – 3 min until embedding medium is completely frozen (opaque white).

NOTE: For TAs, the thickest area (belly) should be visible when on the chuck.

5.5. Place cryostat chuck with the tissue block into the microtome. Uncover blade and advance the microtome forward until just coming into contact with the blade. Section through block at 25 μm sections until the tissue is no longer obscured by embedding medium.

NOTE: While sectioning, adjust the angle of the microtome and/or position of the stage such that the sections are of uniform thickness. It may be helpful to collect a few sections to ensure uniformity in section thickness.

NOTE: It is recommended that the user finds the proper anti-roll plate position prior to sectioning through to the region of interest within the TA (belly), as this allows the user to try several positions of the anti-roll plate until the sections are cut smoothly and do not roll without wasting tissue.

5.6. Change section thickness to 10-12 μm and collect sections on labeled microscope slides. We recommend serial sectioning by collecting adjacent sections on 6-10 slides (labeled 1-x), which allows to stain for multiple markers. If needed, use a thin brush to uncurl sections before collecting on slide.

NOTE: If sections are curling, check that the temperature is holding steady within the −22 to −24 °C range. If there are vertical striations in sections, this could be due to a nick in the anti-roll plate or the blade; this can be fixed by adjusting the position of the anti-roll plate and/or switching to a new blade.

5.7. After collecting adjacent sections of the same sectioning plane onto each slide, adjust thickness back to 25 μm to advance 150-200 μm through the block, then adjust thickness back to 10 μm and begin sectioning again. This serial sectioning allows the user to visualize, image, and quantify at different depths through the TA; our lab tends to use 3-4 serial sections per slide.

5.9. Store slides and tissue blocks at −80 °C.

**6. Immunofluorescent (IF) Staining of Tissue Sections**

NOTE: As antibody concentrations can vary between lots and manufacturers, optimization is recommended by testing several different concentrations of the antibodies on test slides prior to staining the slides of interest.

6.1. Thaw/dry slides either at room temperature or on a warm plate at 37 °C for 10-20 min.

6.2. Use a hydrophobic pen to draw a line at the edge of the slide’s paper surface, where it meets the glass.

6.3. Put the slides in a Coplin jar and wash with 1x PBS + 0.1 % Tween20 (PBST) 3-5 times on a shaker for at least 5 min per wash to rehydrate the tissue sections.

NOTE: At this point, it is important to not let the slides sit without being submerged in PBST (up to hydrophobic line) or the tissue sections will dry out.

6.4. Lay the slides on the rack of a humidifying chamber and overlay the slides with 310-350 μL of blocking solution for 1-2 h at room temperature.

6.5. Prepare the primary antibodies to be used in blocking solution shortly before proceeding to next step.

6.5.1. Primary antibodies for adipocyte staining and whole section imaging: rabbit anti-perilipin (1:1000).

6.5.2. Primary antibodies for lineage tracing of adipocytes: chicken anti-GFP (1:1000) and rabbit anti-perilipin (1:1000)

6.5.3. Primary antibodies for lineage tracing of FAPs: chicken anti-GFP (1:1000) and goat anti-PDGFRα (1:250)

NOTE: It is recommended, when using primary antibodies for the first time, that the user include a negative control slide, in which primary antibodies are not used. This will control for specificity of the antibodies. All other steps in the protocol are followed, including addition of secondary antibodies, but blocking solution is used alone in the next step instead of primary antibody in blocking solution.

6.6. Dump the blocking solution off the slides and overlay with 310-350 μL of blocking solution with primary antibodies and incubate overnight at 4 °C in the humidifying chamber with the lid on.

6.7. Next day, dump the blocking solution/primary antibodies off the slides and place in a Coplin jar. Rinse the slides 2-3 times with PBST and wash 3-5 times with PBST on a shaker for at least 5 min per wash.

6.8. During the last wash, prepare the secondary antibodies or any direct conjugates to be used in blocking solution. Minimize the amount of time these antibodies spend in the light to avoid photobleaching.

6.8.1. Secondary antibodies/direct conjugates: Alexa Fluor 488 donkey anti-chicken (1:1000) or anti-rabbit (1:1000), Alexa Fluor 568 donkey anti-goat (1:1000) or anti-rabbit (1:1000), Alexa Fluor 568 Phalloidin (myofibers; 1:100), DAPI stain (nuclei; 1:500).

6.9. Overlay the slides with 310-350 μL of the blocking solution with secondary antibodies and/or direct conjugates in the humidifying chamber and put the lid on. Incubate at room temperature for 1-2 h. Protect samples from light from now on.

6.10. Dump the blocking solution/secondary antibodies and place in a Coplin jar. Rinse with PBST once and wash 3-5 times with PBST on a shaker for at least 5 min per wash. Keep the Coplin jar covered by a box or other object that does not allow light through to prevent photobleaching.

6.11. Dry the slides as well as possible by tapping the edges and wiping the back against a paper towel, but do not let the tissue sections dry out. Add 3-4 drops of mounting medium to the top horizontal edge of the slide and gently put on a coverslip. Do not press down or move if air bubbles form under the coverslip, any pressure or movement can affect the integrity of the adipocytes.

6.12. Allow mounting medium to set in the dark overnight before imaging.

**7. Whole Mount Immunofluorescent Staining**

7.1. After ~1 h of fixation (see step 3.16. from above), use sharp tipped tweezers to peel away myofibers from the fixed TA.

7.2. Place the separated fibers into a 24-well plate and wash 3×3 min with PBST. For all subsequent incubations, ensure that lid is added to prevent evaporation.

7.3. Incubate for 1 h in permeabilization solution at room temperature (200-300 μl) on shaker, which will allow better penetration of antibodies.

7.4. After rinsing a few times with PBST, overlay with blocking solution (200-300 μl) and block on nutator or shaker overnight at 4 °C.

7.5. Dilute primary antibodies at desired concentration (we find that doubling the concentration is a good starting point) in blocking solution. Incubate samples (200-300 μl) on nutator or shaker overnight at 4 °C.

7.6. Wash samples rigorously with PBST throughout the day with frequent changes at room temperature on shaker, about 4-6 times for 30 min – 1 h each wash.

7.7. Dilute secondary antibodies in blocking solution at desired concentration (1:500 for Alexa secondaries works well) plus nuclear staining and incubate samples (200-300 μl) on nutator or shaker overnight at 4 °C.

7.8. Wash samples rigorously with PBST throughout the day with frequent changes at room temperature on shaker, about 4-6 times for 30 min - 1 h each wash. You can also wash overnight at 4 °C.

7.9. To mount, gently dry off excess PBST and then place the fibers in 1-2 drops of mounting medium on glass slide. To raise the coverslip (18×18mm), add little clay feet. That will prevent the fibers from being squashed and secure the cover slip to the slide. Modeling compounds such as PlayDoh work well for this. Once coverslip is secured, add more medium to the edge until the area under the coverslip is full.

NOTE: Instead of using mounting medium containing anti-fading agents, the tissue can also be moved through an ascending series of Glycerol (30 to 80% Glycerol in PBS).

7.10. Wait for 1-2 days before imaging.

**8. Imaging of intramuscular fat**

8.1. Turn on microscope and launch the imaging software. Secure the slide on the stage.

NOTE: For imaging adipocytes in muscle sections, a 5X or 10X objective combined with widefield microscopy is often sufficient. For visualizing WM-IF, a confocal microscope is required.

8.2. Use ***(GFP / phalloidin / perilipin / DAPI)*** channels to identify the area to be imaged.

8.3. In the imaging software, adjust gain and exposure time for each channel.

8.4. Take images of the whole tissue in each channel (automatic or manual according to microscope and software used) and merge individual tiles to make a composite of the full TA cross section.

NOTE: If the user does not plan to use RT-qPCR to confirm results in adipocyte quantifications, it is recommended they take images of 2-3 different sections of the same TA at different depths. By quantifying the adipocytes in each section and then reporting the average, localized differences in the amount of intramuscular fat due to, for example, injection errors, will be avoided.

**9. Quantifying adipocytes**

9.1. If not previously installed, add the “Cell Counter” plug-in to ImageJ (https://imagej.nih.gov/ij/plugins/cell-counter.html).

9.2. Import images into ImageJ as tif files or original microscope files, such as lif for Leica microscopes. Each channel should be viewed in ImageJ as a separate tif file.

NOTE: If using lif or similar file types, under Bio-Formats Import Options, choose “Hyperstack” for “View stack with:” and check box for “Split channels.” Click “OK” to open file. Also ensure that the “Autoscale” box is unchecked.

9.3. Ensure that images for each channel are in 8-bit format (and gray): Image > Type > 8-bit.

9.4. Merge DAPI (blue), GFP (green), PERILIPIN (red), and PHALLOIDIN (gray) images: Image> Color > Merge channels.

9.5. Check that scale (Analyze > Set Scale) is in microns. Using the freehand selection tool, outline the injured portion of the muscle, then measure and record injured area in excel: Analyze > Measure.

NOTE: Injured muscle can be identified by areas devoid of fibers or with centrally-located nuclei.

9.6. Launch Cell Counter: Plugins > CellCounter > Initialize.

9.7. Select a counter type, then count each adipocyte. Record total number of adipocytes in excel, then calculate the number of adipocytes per 1 mm^2^ of injured area. We use GraphPad/Prism to visualize the data.

**10. Adipogenic gene expression analysis using RT-qPCR**

**10.1. RNA Isolation**

10.1.1. Before beginning, preheat RNase-free water to 45 °C and prepare fresh 70% EtOH (350 μl per sample).

10.1.2 Add 1000 μl of Trizol to each tube containing the sample (see step 3.17. from above). It is important that bead beater-approved tubes are being used.

CAUTION: Trizol is toxic. Wear appropriate personal protective equipment and handle in a fume hood.

10.1.3. Add three medium beads or one large and one small bead to each tube.

10.1.4. Homogenize tissue at 50 Hz for 3 to 5 min using a bead beater. Depending on tissue type and sample size, it may take up to 10 min.

10.1.5. Add 200 μl of chloroform.

CAUTION: Chloroform is toxic. Wear personal protective equipment and handle in a fume hood.

10.1.6. Shake the samples for 15 s.

10.1.7. Incubate for 2 – 3 min at room temperature.

10.1.8. Centrifuge for 15 min at 12,000 g. During this time, add 350 μl fresh 70% ethanol (prepared in step 10.1.1.) to a new Eppendorf.

10.1.9. Pipette out 350 μl of the clear supernatant (upper layer) and add to 350 μl of ethanol in the Eppendorf. Be careful to not aspirate the lower protein and/or DNA layers.

NOTE: Unless RNA is being used for RNAseq experiments, we omit the DNase treatment step as we find that by carefully taking off just the upper 350 μl of the RNA layer, DNA contamination is absent.

10.1.10. Transfer up to 700 μl of the mixture to a RNeasy spin column placed in a 2 mL collection tube. Continue with RNA isolation following the manufacturer’s instructions.

10.1.11. Elute with 30 - 50 μl RNase-free water, depending on expected yield. Keep RNA on ice.

10.1.12. Keep RNA stored at −80 °C.

**10.2. cDNA Synthesis**

10.2.1. Use up to 1 μg of RNA to synthesize cDNA with a cDNA Synthesis Kit, following manufacturer’s instructions.

10.2.2. After the run is complete, add 80 μl of RNase-free water. Store samples at −20 °C.

**10.3. RT-qPCR of Adipocyte-selective Genes**

10.3.1. Using a 384 well format, add 1 μl of primer (~1 μM final concentration) to bottom of each well. Pre-drying primers results in tighter technical replicates. Leave covered until primers have evaporated completely (to speed up evaporation, plate can be placed on heating block set to 37°C).

10.3.2. Set up sample reactions with 4-8 technical replicates as follows: 2.5 μl SYBR Green, 2.1 μl milliQ or RNase-free water, and 0.4 μl cDNA (~1 ng) with 5 μl total volume per well per manufacturer instructions.

10.3.3. Normalize raw CT (cycle threshold) values to housekeeping genes (i.e. *Hprt* and *Pde12*) levels by calculating ΔΔCT as described here ^30^. We use GraphPad/Prism to visualize the data. See ^31^ for primer sequences.

NOTE: Standard practice for RT-qPCR analysis should be followed such as use of a minus Reverse Transcription (-RT) control, PCR reaction and primer validation.

## REPRESENTATIVE RESULTS

### Immunofluorescent Visualization of Intramuscular Fat

Following the steps above, TA tissue sections were gathered from a 21-day post Glycerol injury that were either snap frozen immediately after harvesting or were fixed in 4% PFA for 2.5 h. After cryosectioning and staining, images were taken at the midbelly, the largest area of the TA, using a Leica SPE confocal. As shown in **Figure 1**, PERILIPIN^+^ adipocytes from the fixed TAs have significantly better-preserved morphology compared to unfixed sections, making their identification, visualization, and subsequent quantification much easier and more accurate. To note, we start to detect the first PERILIPIN^+^ lipid droplets at around 5 days post injury, with most adipocytes having formed by day 7. By 21 days post injury, adipocytes have fully matured.

As the amount of fat per TA strongly correlates with the severity of the induced injury, the TAs must be injured significantly to effectively observe and study intramuscular fat formation. We found that practicing injections using ink into cadaver TAs is a great way to improve injury severity. Successful injuries tend to be above 50% of the muscle. To note, injured areas of the muscle represent areas devoid of muscle fibers or areas that are populated by muscle fibers that contain at least one centrally-located nucleus, a known hallmark of a regenerating muscle fiber. We can also readily adopt this protocol to stain for FAPs and fat in 3D. For this, we carefully separate multiple myofibers from the TA post fixation followed by Whole Mount immunofluorescence. The key is to properly secure the fibers to the glass slide and, at the same time, to avoid over-compression of the tissue. By using moldable clay feet, we can adjust the required thickness and secure the cover slip to the slide even allowing the use of an inverted microscope (**Figure 4A**). We have successfully used this method to label PDGFRα^+^ FAPs, Phalloidin^+^ myofibers and PERILIPIN-expressing adipocytes (**Figure 4B**). After obtaining images at multiple z-planes spanning up to 150 μm in thickness, we used the 3D rendering module within the Leica LASX software to create a 3D reconstruction.

**Figure 4.**
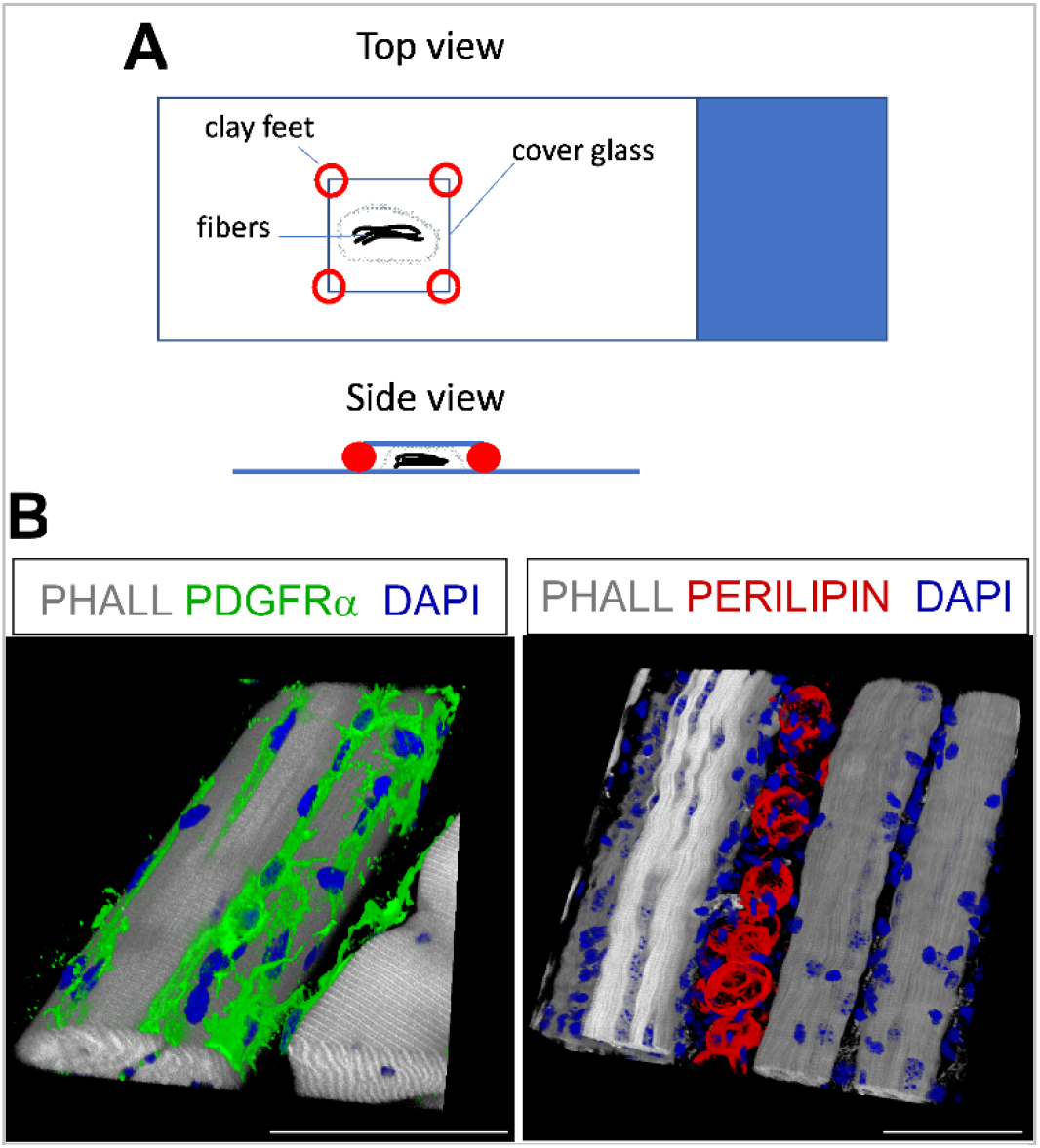
Whole-mount immunofluorescent staining. (A) Top and side view of how to mount the sample and add coverslip for whole-mount staining. (B) Representative 3D reconstructions of FAPs (green) and adipocytes (red) along with myofibers (gray) and nuclei (blue). Scale bars: 50 μm.

### Quantification of intramuscular fat

Once images have been taken of intramuscular fat, we use the Cell Counter function in ImageJ/FIJI to manually count the number of PERLIPIN^+^ adipocytes (**Figure 5A**). Next, we determine the total area of the muscle section as well as the injured area defined by centrally-located nuclei. To control for injury severity, we divide the total number of adipocytes by the injured area resulting in the number of fat cells per 1 mm^2^ of injured muscle. We usually exclude TAs from our quantifications which display <30% injury. As highlighted in **Figure 5B**, a Glycerol injury causes massive amounts of intramuscular fat compared to an uninjured TA muscle. Alternatively, as Perilipin staining is very clean with a high signal to noise ratio, it is also possible to use the “Analyze Particle” function to determine the total area occupied by Perilipin. However, this method will not be able to distinguish between smaller vs. fewer adipocytes. To note, if we are not confirming the amount of intramuscular fat by RT-qPCR (see below), we image and quantify up to 3 sections and report the average number of fat cells present.

**Figure 5.**
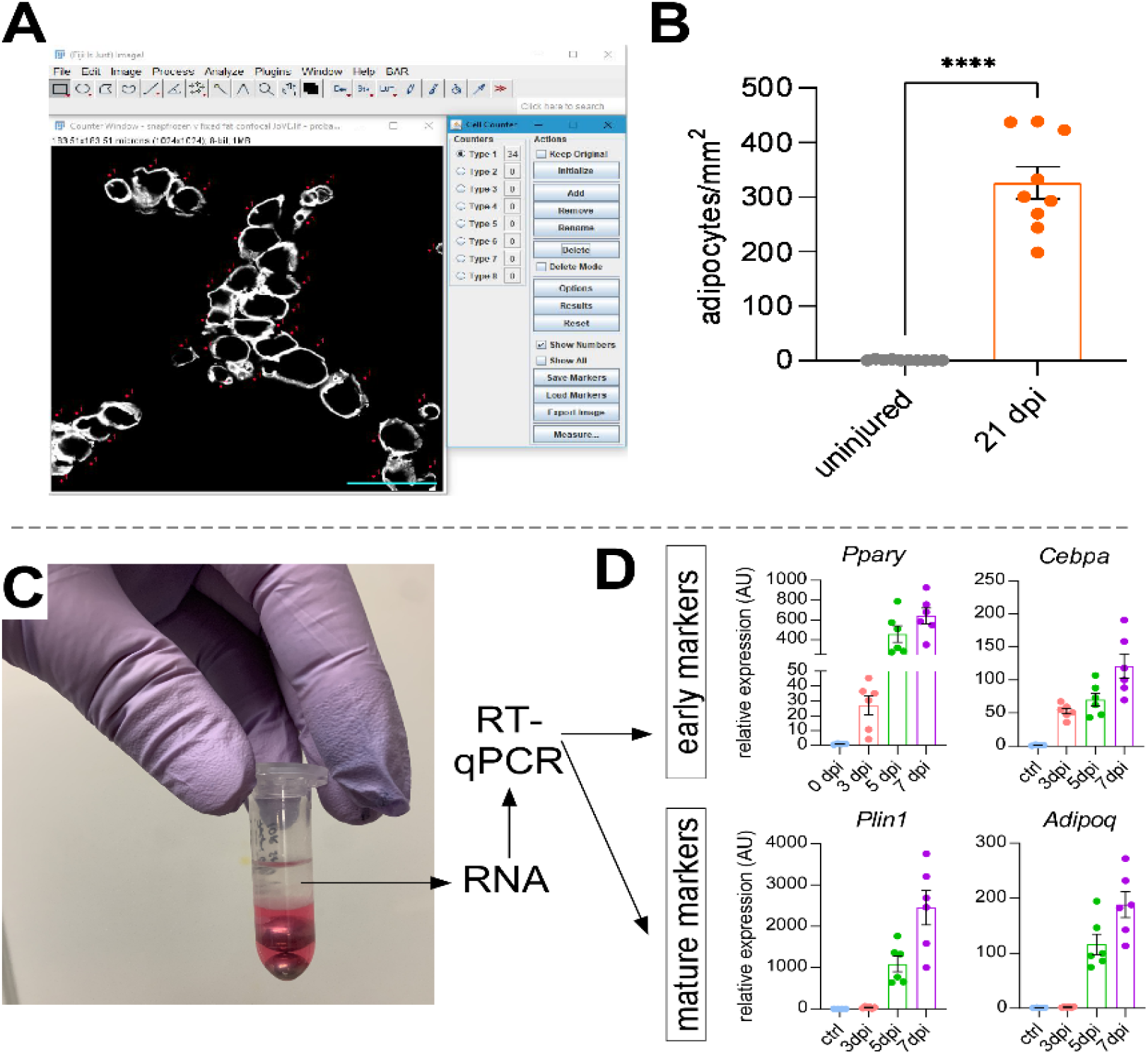
Quantifications of intramuscular fat. (A) Representative image of how to count PERILIPIN^+^ adipocytes (white) using the Cell Counter function in ImageJ; scale bar = 50 μm. (B) Whole TA adipocyte quantifications 21 days post Glycerol injection normalized to 1 mm^2^ of injured area. Error bars shown as SEM. **** = p < 0.0001. (C) RNA layer after homogenization and subsequent phase separation by chloroform is being used for RT-qPCR analysis. (D) Fold changes in expression levels of Pparγ and Cepbα, early adipogenic genes, and Plin1 and Adipoq, two mature adipocyte markers, at different time points post Glycerol injury. Error bars shown as SEM.

To independently confirm the amount of intramuscular fat present, the expression levels of adipogenic markers can be determined. For this, RNA can be isolated from a portion of the same TA muscle used for immunofluorescence (see steps above) at different points post injury. We use a bead beater in combination with Trizol to homogenize the tissue. After adding Chloroform followed by centrifugation, we carefully extract the upper RNA-containing layer and use the Qiagen RNeasy kit for RNA cleanup (**Figure 5C**). This method routinely produces high quality and quantity of RNA suitable for all downstream analyses such as RT-qPCR and RNAseq. For RT-qPCR, we determine the relative expression levels of adipogenic to housekeeping genes using a QuantStudi 6 Flex Real-Time 384-well PCR System and assess any relative changes following the ΔΔCT method ^30^. As described in **Figure 5D**, compared to uninjured TA muscle, Glycerol injury induces expression of early adipogenic markers such as *Pparg* and *Cebpα* as soon as 3 days post injury. Mature markers such as *Adiponectin (Adipoq)* and *Perilipin (Plin1)* can be detected as early as 5 days after Glycerol injury.

### Genetic Lineage Tracing of Adipocytes

Our adipocyte staining protocol can be easily adapted to include genetic lineage tracing of FAPs to map and follow their fate into adipocytes. We have, for example, previously demonstrated that recombination can be induced via tamoxifen administration in *Pdgfrα^CreERT2^; Rosa26^EYFP^*mice 2 weeks prior to injury, effectively removing the floxed stop coding and indelibly activating EYFP expression in FAPs (**Figure 6** and ^31^). We achieve high recombination efficiencies with our tamoxifen regimen with ~75% of PDGFRα^+^ FAPs expressing EYFP ^31^, similar to what other laboratories have reported ^24,32,33^. Demonstrating that FAPs are indeed the cellular origin of intramuscular fat, the majority of FAPs have turned into EYFP^+^ PERILIPIN-expressing adipocytes 7 days post Glycerol injury (**Figure 6** and ^31^).

**Figure 6.**
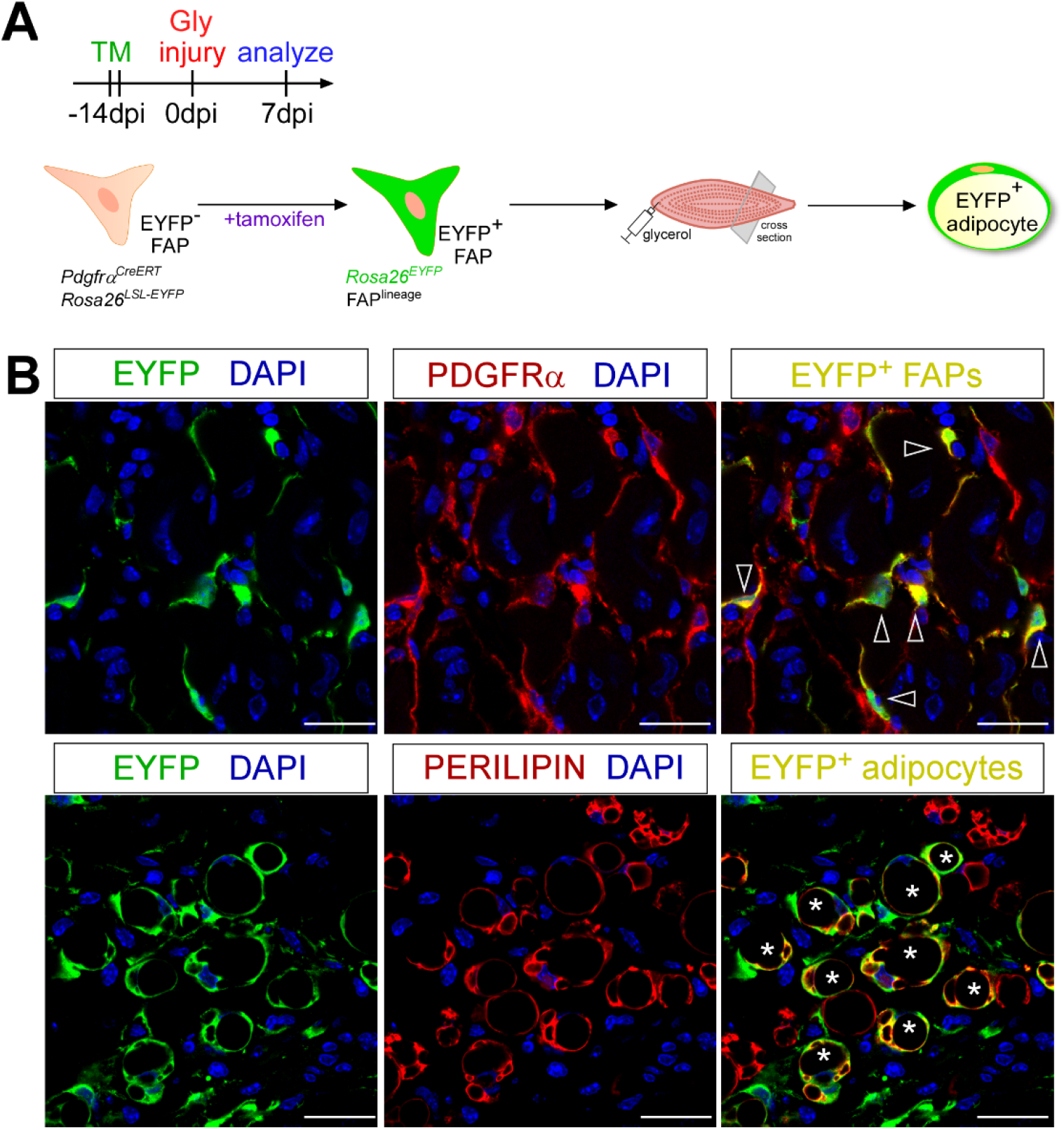
Lineage tracing of FAPs. (A) Schematic overview of experimental setup. (B) Representative immunofluorescent images showing successful recombination and activation of EYFP (yellow) within PDGFRα^+^ FAPs (red, arrowheads) and PERILIPIN^+^ adipocytes (red, asterisks). Scale bars: 25 μm. Figure adapted from ^31^.

## DISCUSSION

Here, we outline an extensive and detailed protocol that allows for efficient visualization and rigorous quantification of intramuscular fat. We also describe how this protocol can be combined with genetic lineage tracing of FAPs to study their conversion into adipocytes under certain conditions.

The most commonly used ways to visualize intramuscular fat are paraffin sections followed by hematoxylin and eosin staining or frozen sections stained for lipophilic dyes such as Oil Red O (ORO). However, while paraffin-processed tissues maintain the best histology, the same process also extracts all lipids preventing the use of lipophilic dyes. Lipophilic staining methods will work on both fixed and unfixed tissue sections. However, lipid droplets are easily displaced by applying pressure to the cover slip, thereby distorting the spatial distribution of intramuscular fat. To circumvent this, a recent study established a rigorous protocol to visualize ORO^+^ adipocytes using a whole mount approach. For this, the authors decellularized the TA allowing them to visualization the spatial distribution of intramuscular fat throughout the whole TA ^34^. However, this technique also prevents the use of other co-stains to mark additional cellular structures. Our whole mount immunofluorescence approach can be used to co-stain adipocytes with a variety of markers allowing for fine mapping of the cellular environment. One major challenge, however, is tissue penetration of the antibodies. The more fibers are kept together, the more difficult it will be for the antibodies to equally penetrate and bind all available antigens. Thus, this method is most effective when looking at small groups of fibers. Similarly, conventional confocal microscopy has a limited focal depth. However, with the current development of novel tissue clearing methods plus new imaging technology, greater tissue penetration and visualization will be possible in the future ^35–37^.

While prior fixation of muscle tissue preserves adipocyte morphology, it also creates a challenge to assess the size of myofibers, an important measurement of muscle health. Myofiber size is determined by measuring the cross-sectional area of myofibers. We have previously reported that prior fixation of muscle tissue will cause most markers available to outline myofibers to fail ^28^. To overcome this hurdle, we have developed a novel image segmentation pipeline, which allows the measurement of myofiber size even in fixed muscle sections ^28^. Thus, we have established a robust and efficient tissue processing pipeline that, combined with this protocol, overcomes most disadvantages caused by prior fixation of muscle tissue.

Another major advantage of our approach is versatility. By splitting the TA into two parts, we are maximizing the amount of information we can obtain from one muscle. This not only reduces animal numbers but also adds an extra layer of control by confirming histology through gene expression and vice versa. In addition, many different genes can be examined beyond adipogenic genes. The isolated RNA can also be used for a whole muscle RNAseq experiment. Finally, the snap frozen muscle piece can also be used for protein work.

Together, this protocol outlines a robust, efficient and rigorous tissue processing pipeline that will allow visualization and quantification of intramuscular fat, the first step in developing novel treatment options to combat fatty fibrosis. At the same time, it is versatile and can be adopted to many different cell types within muscle as well as adipocytes in other tissues.

